# A strategy for complete telomere-to-telomere assembly of ciliate macronuclear genome using ultra-high coverage Nanopore data

**DOI:** 10.1101/2020.01.08.898502

**Authors:** Guangying Wang, Xiaocui Chai, Jing Zhang, Wentao Yang, Chuanqi Jiang, Kai Chen, Wei Miao, Jie Xiong

## Abstract

Ciliates contain two kinds of nuclei: the germline micronucleus (MIC) and the somatic macronucleus (MAC) in a single cell. The MAC usually have fragmented chromosomes. These fragmented chromosomes, capped with telomeres at both ends, could be gene size to several megabases in length among different ciliate species. So far, no telomere-to-telomere assembly of entire MAC genome in ciliate species is finished. Development of the third generation sequencing technologies allows to generate sequencing reads up to megabases in length that could possibly span an entire MAC chromosome. Taking advantage of ultra-long Nanopore reads, we established a simple strategy for the complete assembly of ciliate MAC genomes. Using this strategy, we assembled the complete MAC genomes of two ciliate species *Tetrahymena thermophila* and *Tetrahymena shanghaiensis*, composed of 181 and 214 chromosomes telomere-to-telomere respectively. The established strategy as well as the high-quality genome data will provide a useful approach for ciliate genome assembly, and a valuable community resource for further biological, evolutionary and population genomic studies.

## Introduction

Ciliate separates its germline and somatic genetic information by maintaining two kinds of functionally distinct nuclei: the diploid micronucleus (MIC), and the polyploid macronucleus (MAC) (Gorovsky, 1973; Lynn, 2008). The MIC, like other eukaryotes, usually contains long chromosomes with centromeres and capped by telomeres. In general, the MAC genome comes from the MIC genome through a so-called MAC differentiation process in the sexual stage (conjugation) of ciliate (Orias, 2000). In MAC differentiation, the MIC-like chromosomes are fragmented into small pieces at the chromosome breakage sites (CBSs), and the internal eliminated sequences (IESs), which contain transposable elements, are removed (Orias, 2000). This process finally results in the MAC containing fragmented chromosomes with length range from gene size to several megabases, and capped by telomere sequences at both ends but without centromeres.

Development of the third generation sequencing technologies, e.g. the Nanopore sequencing, allows to generate sequencing reads up to megabases in length (Jain et al., 2018), and thus could sometimes sequence an entire MAC chromosome of ciliate by a single read. The generation of such long sequencing reads gives the opportunity to assemble more complete MAC genomes of ciliates.

Here, we reported a simple strategy which was used to assemble the complete genome of *T. thermophila* and *T. shanghaiensis* using high coverage Nanopore sequencing data.

## Materials and Methods

### Cell culture and DNA extraction

*T. thermophila* SB210 and *T. shanghaiensis* (ATCC accession: 205039) cells were grown in SPP medium (Cassidy-Hanley, 2012) and harvested at a density of 250,000 cells/ml. The total DNA was extracted using the Blood & Cell Culture DNA Midi Kit (Q13343, Qiagen, CA, USA) following the manufacturer’s protocol. The DNA was then purified using the Agencourt AMPure XP beads (A63881, BECKMAN), and the DNA quality and quantity was tested using both NanoDrop One UV-Vis spectrophotometer (Thermo Fisher Scientific, USA) and Qubit 3.0 Fluorometer (Invitrogen, USA).

### Nanopore sequencing

Approximately 10 µg of DNA was size-selected (10-50 Kb) using Blue Pippin (Sage Science, Beverly, MA), and sequencing library was constructed using the Ligation sequencing 1D kit (SQK-LSK108, ONT, UK) according to the manufacturer’s instructions. Each library was sequenced on R9.4 FlowCells using the PromethION sequencer (ONT, UK) for 48 hours. Base calling was subsequently performed on fast5 files using the ONT Albacore software (v0.8.4), and the “passed filter” reads (high quality data) were used for downstream analysis.

### Genome assembling and polishing

Genome assembling was performed using 60X Nanopore datasets. Assemblers, including CANU (Koren et al., 2017), NECAT (https://github.com/xiaochuanle/NECAT), SHASTA (https://github.com/chanzuckerberg/shasta), Flye (Kolmogorov, Yuan, Lin, & Pevzner, 2019), and wtdbg2 (Ruan & Li, 2019), were used. The parameters for the assemblers are listed as follows: 1) CANU, -fast genomeSize=100m; 2) NECAT, GENOME_SIZE=100000000 MIN_READ_LENGTH=3000; 3) SHASTA, default settings; 4) Flye, -g 100m; 5) wtdbg2, default settings. The performance of CANU and NECAT far better than three other assemblers in assembling the MAC chromosomes capped with telomere sequences in both ends. Comparing to CANU, the time cost of NECAT was far less than CANU, and thus NECAT was recommended. Quickmerge (https://github.com/mahulchak/quickmerge) was used to merge the un-closed scaffolds to the 60X genome assemblies (command line: merge_wrapper.py un-closed_scaffolds 60X_assembly). After each round of merging, the closed scaffolds (MAC chromosomes) were extracted, and the left un-closed scaffolds were used to perform the next round of merging. After that, an addition round of merging between the un-closed scaffolds and error corrected telomere-sequences-containing reads was performed using miniasm (−1 -2 -c 1) (Li, 2016). Final genome polishing was performed based on the Illumina sequencing data using Pilon (https://github.com/broadinstitute/pilon).

### Results and Discussion

*T. thermophila* is a very useful unicellular model organism for molecular and cellular biology (Ruehle, Orias, & Pearson, 2016). In 2006, the MAC genome of *T. thermophila* has been sequenced using the Sanger method (Coyne et al., 2008; Eisen et al., 2006), which greatly accelerated the studies using *Tetrahymena* system. The current MAC genome assembly (103.0 Mb, http://ciliate.org/index.php/home/downloads) of *T. thermophila* has 1158 scaffolds, among which 128 (∼58.9 Mb) were capped by telomeres with C4A2 repeats at 5’-end and G4T2 repeats at 3’-end (hereafter defined as closed scaffolds) and could be regarded as complete MAC chromosomes. However, about a half of genome sequences, composed of 1030 scaffolds, are still not assembled as complete MAC chromosomes (hereafter defined as un-closed scaffolds).

About 1000X Nanopore sequencing data (total DNA of both MAC and MIC, reads N50: 25.8 Kb) were obtained to finish the MAC genome assembly. Comparison of different third-generation sequencing data assemblers,including CANU, NECAT, SHASTA, Flye and wtdbg2, were performed. In practice, CANU and NECAT showed better performance on assembling closed scaffolds compared to other assemblers. We divided the ∼1000X Nanopore data into different parts, each with ∼60X data, and individually assembled them (Figure 1). We have two reasons to do this division: 1) The MIC reads (contaminations) could be limited below 3X (the copy number ratio between MAC and MIC is 45:2), which will usually be filtered by genome assemblers (Jain et al., 2018); 2) At 60X coverage, CANU and NECAT already have good assembling performance and the time cost of assembling could be greatly reduced.

**Figure 1.**
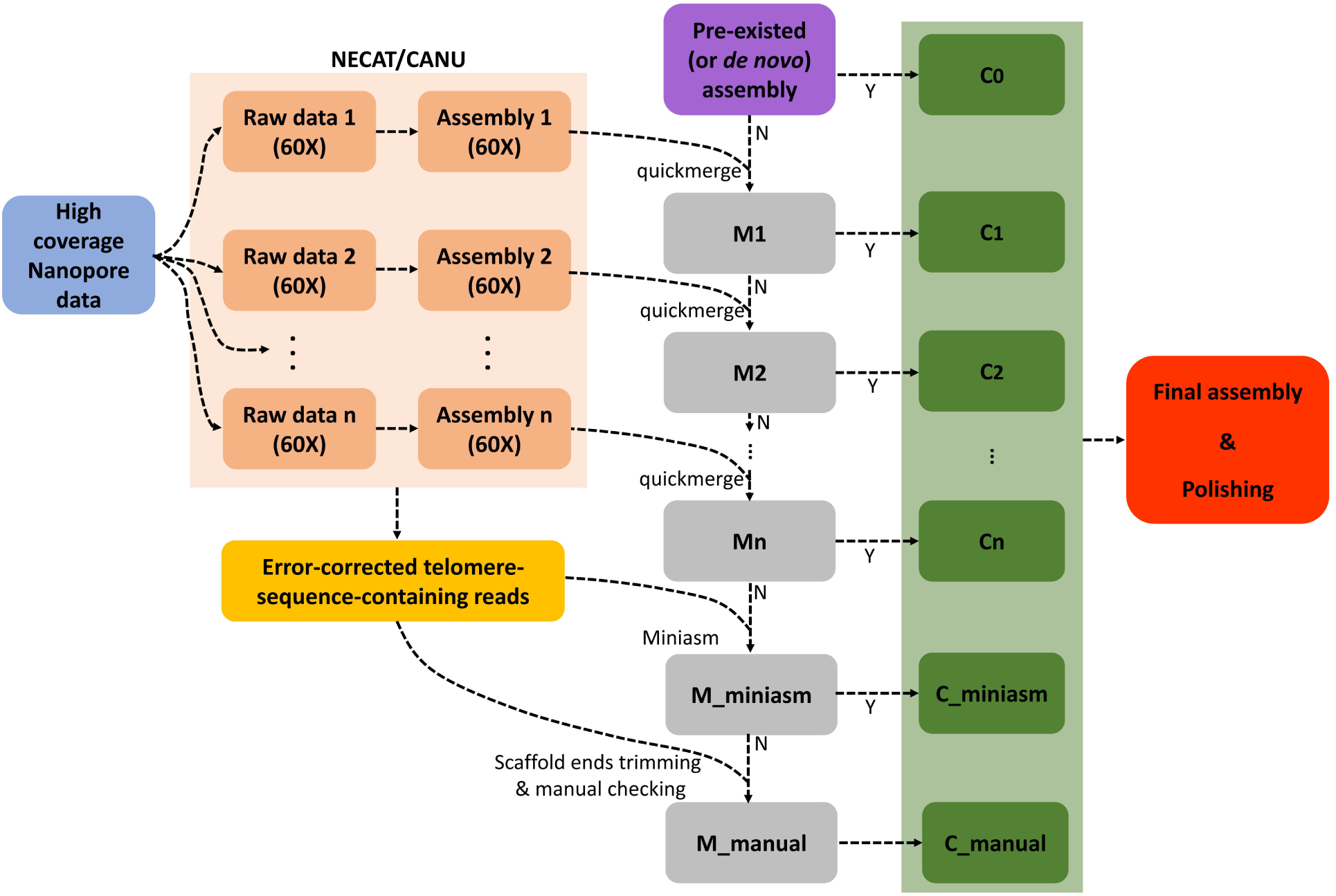
Diagram showing the strategy to assemble complete MAC genome of ciliate. M1 to Mn, the un-closed scaffolds in each round (1 to n) which do not have telomere sequences in both ends. M_miniasm means the un-closed scaffolds after merging using miniasm. C1 to Cn, the closed scaffolds (MAC chromosomes) in each round (1 to n) which have telomere sequences in both ends. C_miniasm means the closed scaffolds (MAC chromosomes) after merging using miniasm. C_manual means the closed scaffolds after the manual checking of overlaps between TSCR and un-closed scaffolds (trimmed).

We started from the 1158 scaffolds in current genome assembly of *T. thermophila* (Figure 1), and divided these scaffolds into two parts: 1) 128 closed scaffolds which assembled as complete MAC chromosomes; 2) 1030 un-closed scaffolds which have not been assembled as MAC chromosomes. For the 128 closed scaffolds, three of them still have gaps (one per each). These gaps were easily closed by aligning the three scaffolds to the 60X Nanopore data assemblies. The left 1030 un-closed scaffolds were iteratively merged with each assembled genome using 60X Nanopore data (Figure 1). After six rounds of merging using quickmerge, 34 closed scaffolds were newly obtained. After that, we extracted the 256,181 raw telomere-sequence-containing reads (TSCR, reads N50, 28.5 Kb) from Nanopore data (Figure 1), and sequencing errors were corrected using NECAT. These error corrected TSCR were aligned to the left scaffolds using minimap2, and followed by a new round of assembly using miniasm (Figure 1), and additional 12 scaffolds with telomere sequences capped at both ends were obtained, and only six scaffolds (3.3 Mb) could not be resolved. To close these six scaffolds, we manually checked the overlaps between TSCR and these scaffolds (Figure 1), and all of them were closed by trimming their terminal sequences and re-merging with TSCR.

In summary, the complete MAC genome (102.9 Mb) with a total of 181 MAC chromosomes (including rDNA mini-chromosome) were obtained. These MAC chromosomes were re-named from 1 to 181 by their order along the five MIC chromosomes. Figure 2 showed the full panel of the 181 MAC chromosomes. The longest MAC chromosome is 3.26 Mb in length, and the shortest one (excluding rDNA mini-chromosome) is 38 Kb in length. The real N50 of the MAC genome is ∼891 Kb. A total of 22 classes of repetitive sequences, which masked 5.2% MAC genome, were identified by RepeatModeler. The repetitive sequences in the MAC are not randomly distributed, most of them are enriched in the MAC chromosomes and derived from the pericentromeric and subtelomeric regions of MIC chromosomes (Hamilton et al., 2016; Xiong et al., 2019). In particular, we also found some new genes which missed in the current genome assembly, for example, the alpha 2 subunit of the proteasome.

**Figure 2.**
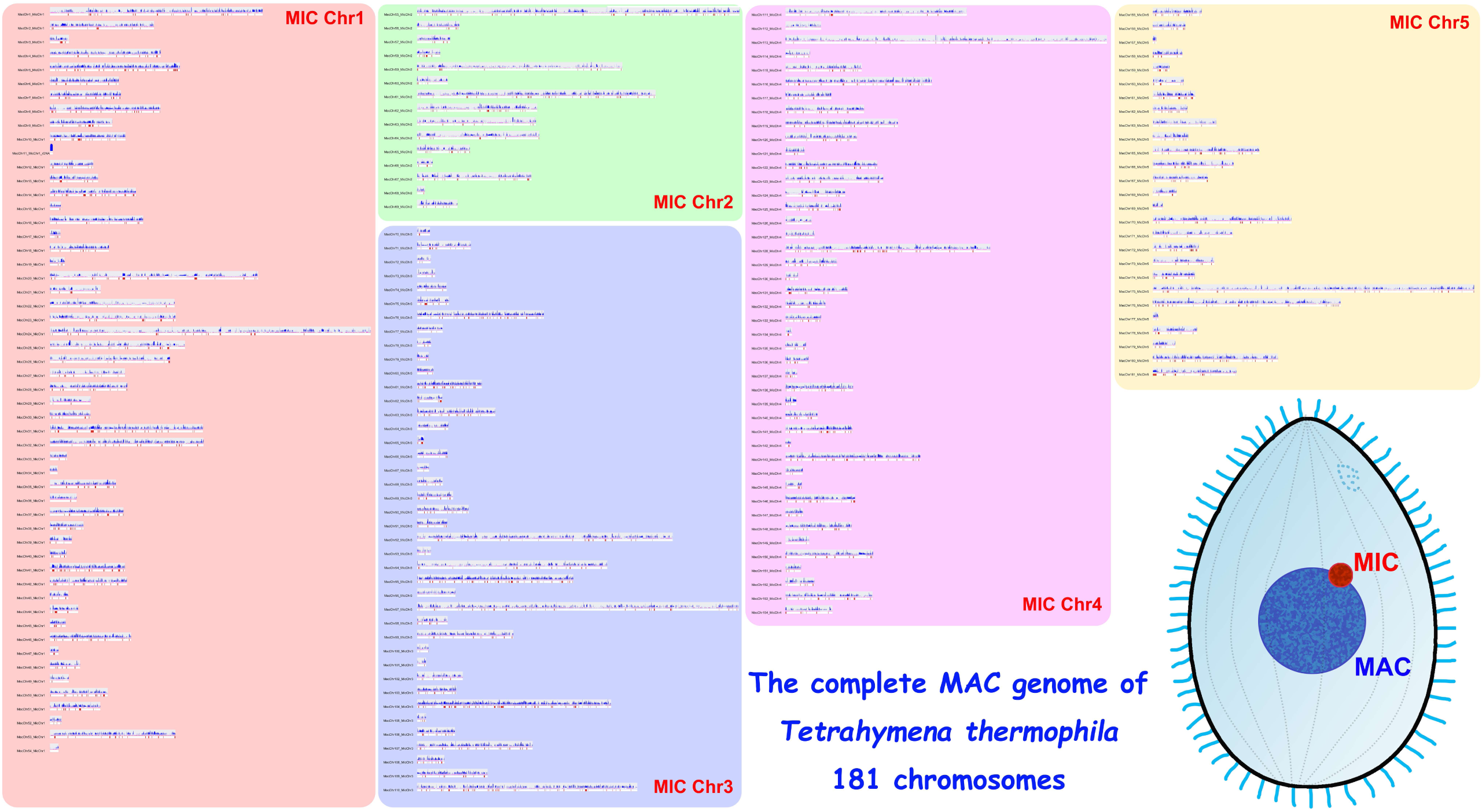
A full panel of 181 MAC chromosomes of *T. thermophila*. For each MAC chromosome, the pink boxes represents the predicted genes; the red boxes represent all the genes that have been named in TGD wiki (http://ciliate.org/); the blue histogram represents the gene expression profile across the chromosome in vegetative growth (Xiong et al., 2012).

To test the applicability of this strategy, we generated ∼900X Nanopore sequencing data (reads N50: 30.8 Kb) of *T. shanghaiensis*. Instead using pre-existed assembly, we started from a 60X *de novo* assembly by NECAT, and then followed the strategy showing in Figure 1. After eight rounds of merging using quickmerge and a round of assembly using miniasm, and followed by additional manual checking, we finally got the complete genome of *T. shanghaiensis* with 214 MAC chromosomes (92.0 Mb) which capped with telomere sequences at both ends. Genome assembly statistics of the two *Tetrahymena* species are shown in Table 1. We anticipate that the established strategy can probably be used directly or with a slight adaptation to assemble complete MAC genomes of other ciliate species.

**Table 1.**
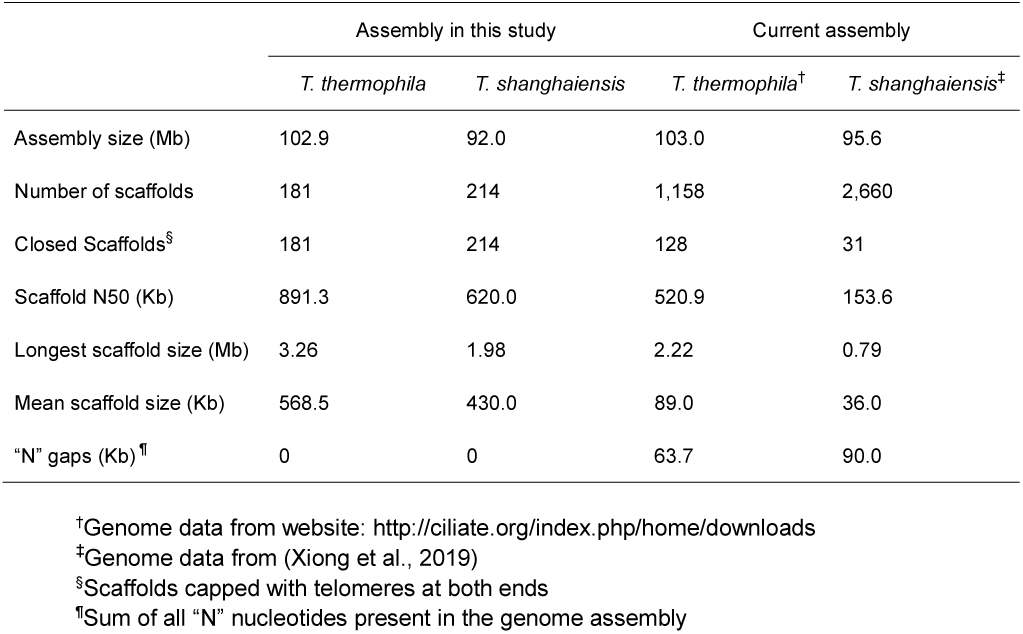
Genome assembly statistics of *T. thermophila* and *T. shanghaiensis*

## Acknowledgments

This work was supported by the National Natural Science Foundation of China Grant no. 31525021, 91631303 (to W.M.), 31672281, 31872221 (to J.X.) and 31900316 (to G.W.); Knowledge Innovation Program of the Chinese Academy of Sciences (to J.X.); Youth Innovation Promotion Association, Chinese Academy of Sciences (to J.X.); The bioinformatics analysis was aided by the computing resources of the Wuhan Branch, Supercomputing Center, Chinese Academy of Sciences, China.

## Author contributions

W.M. and J.X. designed the project. J.X., G.W. and W.Y. assembled and annotated the genome. X.C., J.Z., C.J. and K.C. prepared DNA samples for sequencing. J.X. and G.W. wrote the manuscript. All authors read, revised and approved the final manuscript.

## Data accessibility

The complete genome sequences of *T. thermophila* and *T. shanghaiensis* can be accessed from http://ciliate.ihb.ac.cn/tcgd/download.html.

